# Aerosol Exposure of Cynomolgus Macaques to SARS-CoV-2 Results in More Severe Pathology than Existing Models

**DOI:** 10.1101/2021.04.27.441510

**Authors:** Sandra L. Bixler, Christopher P. Stefan, Alexandra Jay, Franco Rossi, Keersten M. Ricks, Charles J. Shoemaker, Alicia M. Moreau, Xiankun Zeng, Jay W. Hooper, David Dyer, Ondraya Frick, Jeffrey W. Koehler, Brian Kearney, Nina DiPinto, Jun Liu, Samantha Tostenson, Tamara L. Clements, Jeffrey M. Smith, Joshua A. Johnson, Kerry Berrier, Heather Esham, Korey L. Delp, Susan R. Coyne, Holly Bloomfield, Paul Kuehnert, Kristen Akers, Kathleen Gibson, Timothy D. Minogue, Aysegul Nalca, Margaret L. M. Pitt

## Abstract

The emergence of SARS-CoV-2 pandemic has highlighted the need for animal models that faithfully recapitulate the salient features of COVID-19 disease in humans; these models are necessary for the rapid down-selection, testing, and evaluation of medical countermeasures. Here we performed a direct comparison of two distinct routes of SARS-CoV-2 exposure, combined intratracheal/intranasal and small particle aerosol, in two nonhuman primate species: rhesus and cynomolgus macaques. While all four experimental groups displayed very few outward clinical signs, evidence of mild to moderate respiratory disease was present on radiographs and at the time of necropsy. Cynomolgus macaques exposed via the aerosol route also developed the most consistent fever responses and had the most severe respiratory disease and pathology. This study demonstrates that while all four models were suitable representations of mild COVID-like illness, aerosol exposure of cynomolgus macaques to SARS-CoV-2 produced the most severe disease, which may provide additional clinical endpoints for evaluating therapeutics and vaccines.

## Introduction

The threat of a previously unknown emerging pathogen has long been a concern of the scientific and medical communities. This includes the challenge of rapidly developing and implementing scientific tools for characterization and investigation of the new threat, as well as the production and deployment of vaccines and therapeutics. Although the discovery of severe acute respiratory syndrome (SARS-CoV) and Middle East respiratory syndrome (MERS-CoV), both caused by coronaviruses, have demonstrated this need on a limited scale, the emergence of COVID-19 from Wuhan, China in late 2019 and the subsequent identification of SARS-CoV-2 as the causative agent, has provided an example of this process on an unprecedented global scale. The key components of a “toolbox” required for responding to any emerging pathogen includes characterized agent for use as reference material, reagents such as antigen and antibody for serological assays, assays for rapid detection of infected individuals, and animal models for evaluation of pathogenesis and medical countermeasures, among others.

Although three vaccines for SARS-CoV-2 have already received emergency use authorization (EUA) from the Food and Drug Administration (FDA) to-date, with additional vaccine candidates currently being evaluated in clinical trials, the development of relevant animal models for COVID-19 remains an unmet need. Animal models that faithfully replicate the salient aspects of human COVID-19 disease are not only crucial for down-selecting early-stage medical countermeasures, but also for understanding the pathophysiological changes and immunological processes resulting from COVID-19 disease. While a number of animal models for SARS-CoV-2 have been developed including genetically-modified mice, hamsters, and nonhuman primates (NHPs) (1), additional development and refinement for SARS-CoV-2 animal models is still needed. Moreover, although antibody responses are likely involved, the correlate of protection for SARS-CoV-2 still remains unknown (2). The NHP model, which is commonly considered a “gold standard” in infectious disease research due to its close fidelity to human disease and immunology, will be important for bridging potential correlates of protection to the immune responses generated in humans.

As with other members of the betacoronavirus family, SARS-CoV-2 causes respiratory disease of variable severity in humans, from mild illness resembling the common cold to severe respiratory distress resulting in death (3–6). The clinical signs of COVID-19 most commonly include fever, cough, and shortness of breath, with a subset of patients developing gastrointestinal and neurological signs (3–6). Severe respiratory disease is characterized by pneumonia, “ground-glass” opacity, and consolidation on chest radiographs (3–6) and histopathological findings of alveolar damage, multinucleated giant cells, congestion and hemorrhage, inflammatory infiltrates, and fibrin deposition (7). The development of NHP models of COVID-19 have been largely focused on routes of infection that may reflect and/or simulate potential respiratory and mucosal modes of transmission, such as intranasal (IN), intratracheal (IT), oral, ocular, and combinations thereof (8–13). Generally, most NHP models for SARS-CoV-2 have used rhesus macaques (RM), cynomolgus macaques (CM), or African green monkeys (AGM) and have resulted in very mild respiratory disease with minimal to no mortality (8–13). Severe respiratory distress and mortality have only been observed in instances where older animals were utilized (12). Previous NHP model evaluation performed at the United States Army Medical Research Institute of Infectious Diseases (USAMRIID) has suggested that there are likely species-associated differences in type and severity of clinical disease signs following aerosol (AE) exposure to SARS-CoV-2, with AGMs and CMs demonstrating the most consistent presentation of disease (13).

While a number of potential routes of transmission have been proposed for SARS-CoV- 2, contact, droplet, and aerosol likely represent the primary modes of transmission, in keeping with other respiratory viruses (14–16). Droplets are thought to be produced by coughing, sneezing, and talking and generally are > 5 µm in size, meaning they do not remain suspended in the air over long periods of time and/or distances (17–18). Aerosols are considered to be ≤ 5 µm and can remain suspended in the air for long distances and periods of time (16–17). While the relative contribution of each of these routes to SARS-CoV-2 transmission is still unknown, reports of transmission between individuals separated by greater distances (> 2 m) and in poorly ventilated indoor areas suggests that aerosol transmission is indeed occurring (14, 16). In studies of SARS-CoV-2 and other respiratory pathogens, combined IT/IN administration is often used as a surrogate exposure route to achieve exposure of both the upper and lower respiratory tracts to pathogen. Although the IT/IN is often the more favored route due to the relative ease of performance in a laboratory setting, AE may represent a more natural route of exposure and may be useful in developing an animal model that accurately captures the clinical features of human COVID-19 disease. In addition to prolonged suspension, the generation of small aerosol particles enables deeper penetration into the lungs (17), potentially resulting in more severe respiratory disease and pathology. As the majority of the current NHP models have failed to replicate the more severe cases of disease seen in some COVID-19 patients, AE represents a potential opportunity to fill this critical gap. Additionally, USAMRIID’s AE delivery system incorporates a head-only exposure chamber, ensuring that animals are also exposed to virus through mucosal surfaces including the mouth and eyes which may replicate the multifaceted mode of COVID transmission in humans.

The majority of medical countermeasure studies, including those supported by the federal response, have used the combined IT/IN instillation of SARS-CoV-2 in RMs as the model of choice. To date, there is no published data on the direct comparison of AE to IT/IN for SARS-CoV-2. Here, we performed a head-to-head comparison of these two routes of exposure in two nonhuman primate species: RMs and CMs.

## Results

### SARS-CoV-2 Infection of RM and CM Exposed by AE or IT/IN

A total of eight RMs and eight CMs were exposed to the WA-1/2020 strain of SARS-CoV-2. Four of each species were infected by combined IT/IN administration, while the remaining four animals of each species were exposed to small particle AE. To develop the most stringent model, the goal was to expose the animals to the highest possible dose of virus achievable based on route of exposure and the titer of the virus stock. Based on a titer of 5.45x10^6^ plaque forming units (pfu)/mL, the target dose was 2x10^7^ pfu for the IT/IN exposure route and 5x10^4^ – 5x10^5^ pfu for AE route. The actual dose received by the animals in the IT/IN group as determined by neutral red plaque assay was 2.65x10^7^ pfu. The AE animals received between 4.45x10^4^ – 8.79x10^4^ pfu, with a mean inhaled dose of 5.85x10^4^ pfu for RMs and 6.66x10^4^ pfu for CMs.

A variety of biosamples including blood, nasopharyngeal (NP) swabs, oropharyngeal (OP) swabs, rectal swabs, saliva, and bronchoalveolar lavage (BAL) fluid, were periodically collected from animals to confirm SARS-CoV-2 infection, to monitor infection kinetics and viral replication, and to detect potential shedding from mucosal surfaces (**Figure 1A**). Using quantitative reverse transcription polymerase chain reaction (qRT-PCR), the presence of viral RNA was detected on Day 2 PE in NP and OP swabs from all animals, regardless of species or exposure route (**Figure 1B and Suppl. Figure 1A**). In most instances, peak RNA levels were observed at this time point (7.98-12.47 log10 target copies/mL). The kinetics of viral RNA in NP swabs were generally consistent between the groups, with a gradual decline in viral RNA levels after Day 2 PE and more variability present at the later sampling time points (Days 6 and 8 PE). This was in contrast to OP swabs which showed a more dramatic decrease in viral RNA by Day 4 and a trend towards increased viral titers in RMs (AE and IT/IN) on Day 8 PE as compared to CMs (AE and IT/IN). In rectal swabs, viral RNA was only detected in one CM (CM IT/IN 4) as compared to six RMs (4 from RM IT/IN and 2 from RM AE) on Day 2 PE (**Suppl. Figure 1B**).

**Figure 1.**
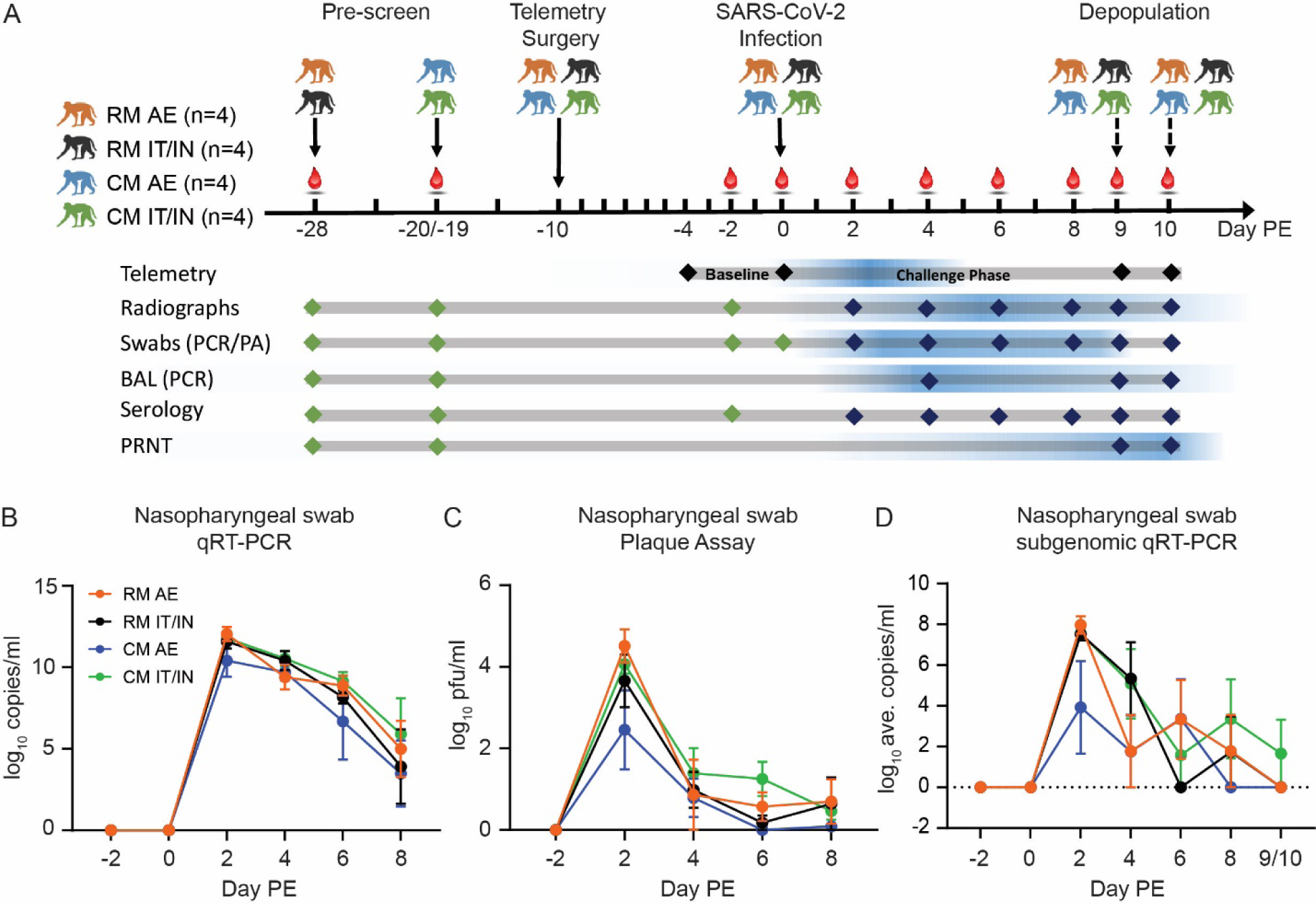
Infection of RM and CM with SARS-CoV-2 by AE or IT/IN exposure. A. Study design. Red droplets denote phlebotomy days. Baseline sampling points are represented by green diamonds, while dark blue represents post-infection sampling points. The blue gradient provides a visual representation of the peak responses observed from each assay/measurement. B-D. Detection of viral RNA by qRT-PCR (B), infectious virus by plaque assay (C), and subgenomic RNA by real-time RT-PCR in NP swabs. Data is shown as the group mean ± SEM.

Interestingly, no viral RNA was detected from any rectal swabs on Day 4 PE, but was detected in the majority of the animals in the RM AE, RM IT/IN, and CM IT/IN groups on Day 6 PE. The absence of detectable viral RNA on Day 4 PE may suggest a possible sampling error. Also of note, viral RNA was never detected in rectal swabs from any of the CM AE animals. Detection of viral RNA in saliva samples was variable between Days 2 and 8 PE, and absent after Day 8 PE (data not shown).

While no viable virus was detected in any rectal swabs by neutral red plaque assay (data not shown), the presence of viable virus was confirmed for both NP and OP swabs. Virus was detected for the majority of animals by Day 2 PE for both NP and OP swabs, which often represented peak titers (**Figure 1C and Suppl. Fig. 1C**). Peak levels in NP swabs were between 3.70-5.34 log10 pfu/mL for RM AE, 1.80-4.78 log10 pfu/mL for RM IT/IN, 1.84-4.19 log10 pfu/mL for CM AE, and 3.40-4.44 log10 pfu/mL for CM IT/IN. Peak levels in OP swabs were between 4.00-4.44 log10 pfu/mL for RM AE, 0.35-3.70 log10 pfu/mL for RM IT/IN, 1.48-3.26 log10 pfu/mL for CM AE, and 2.22-3.74 log10 pfu/mL for CM IT/IN. For both NP and OP swabs, the highest titers were present in the RM AE group. While significant differences between swab types and experimental groups were not observed, a trend was observed in that virus typically persisted in NP swabs longer than in OP swabs for most animals. By Day 8 PE, virus was still detected in only half the animals and more often in NP swabs than OP swabs.

The presence of infectious virus in the plaque assay was supported by the detection of subgenomic RNA, indicative of replicating virus. All animals had detectable subgenomic RNA in either NP swabs or BAL fluid during the study, with the exception of CM AE 3 (**Figure 1D and Suppl. Fig. 1D**). In general, Day 2 PE represented the peak subgenomic titers in NP swabs from all groups, with the CM AE animals having the lowest average titer on this day. This mirrors the trend observed in the plaque assay data. Subgenomic titers decreased from Day 2 PE to Day 8 PE, with some fluctuation from time point to time point. Only one animal (CM IT/IN 1) still had detectable subgenomic RNA at the terminal time point (Days 9/10 PE). Although fewer sampling time points were available for the BAL samples, subgenomic RNA was detectable in a number of animals on Day 4 PE and only one animal (RM AE 3) at the time of euthanasia.

### Early Physiological Changes Following SARS-CoV-2 Infection

All animals were implanted with M00 telemetry devices to monitor body temperature and activity over the course of infection. This enabled the detection of transient changes in temperature that might otherwise be overlooked by periodic, infrequent collection of rectal temperatures during anesthetized physical examination. Fever was the earliest clinical sign that developed following SARS-CoV-2 exposure, appearing as early as 17 hours post-infection (**Figure 2**). All of the CMs in the AE and IT/IN groups developed fever at some point during the course of the study, although fever was not sustained (i.e. each fever episode was shorter than 2 hours) in two of the CM IT/IN animals (CM IT/IN 3 and 4). Only one RM AE animal (1/4; RM AE1. 2) and two RM IT/IN animals (2/4; RM IT/IN 3 and RM IT/IN 4) developed fever during the study. The mean maximum temperature change for each group was 1.4°C for RM AE (range = 0.5-2.8°C), 2.0°C for RM IT/IN (range = 1.1-3.3°C), 2.7°C for CM AE (range = 2.3-3.2°C), and 2.2°C for CM IT/IN (1.7-3.1°C) (**Figure 2B**). Fever hours (Fever-h) is the sum of the significant temperature elevations (defined as >3 standard deviations above baseline) in a 24-hr period; this measure gives an indication of the intensity of the fever by calculating the area between the baseline and the post-exposure temperature curves. The greatest fever-h for each group was 4.6 °C-h for RM AE (range = 1.0-9.6 °C-h), 9.9 °C-h for RM IT/IN (range = 2.8-21.9 °C-h), 15.8 °C-h for CM AE (range = 5.5-25.2 °C-h), and 14.5 °C-h for CM IT/IN (range = 9.5-18.5 °C-h) (**Figure 2A**). The daily percent of significant temperature elevations (TEsig) duration is the % of the 24-hour daily time period where body temperatures were significantly elevated (greater than 3 SD for that time period). The peak percentage of TEsig values were 46% in the CM AE group, 19% in the RM AE group, 45% in CM IT/IN group, and 31% in the RM IT/IN group. Taken together, CM appeared to have a greater fever response than RM in terms of both maximum temperature change and fever-h, with CM AE having the highest values. In terms of route, the IT/IN animals appeared to have the shortest duration of fever, as fever responses were limited to Day 1 PE. In the AE animals, elevated temperatures generally lasted through Day 2 PE, although one CM AE animal (CM AE 4) had a fever response that persisted to Day 4 PE.

**Figure 2.**
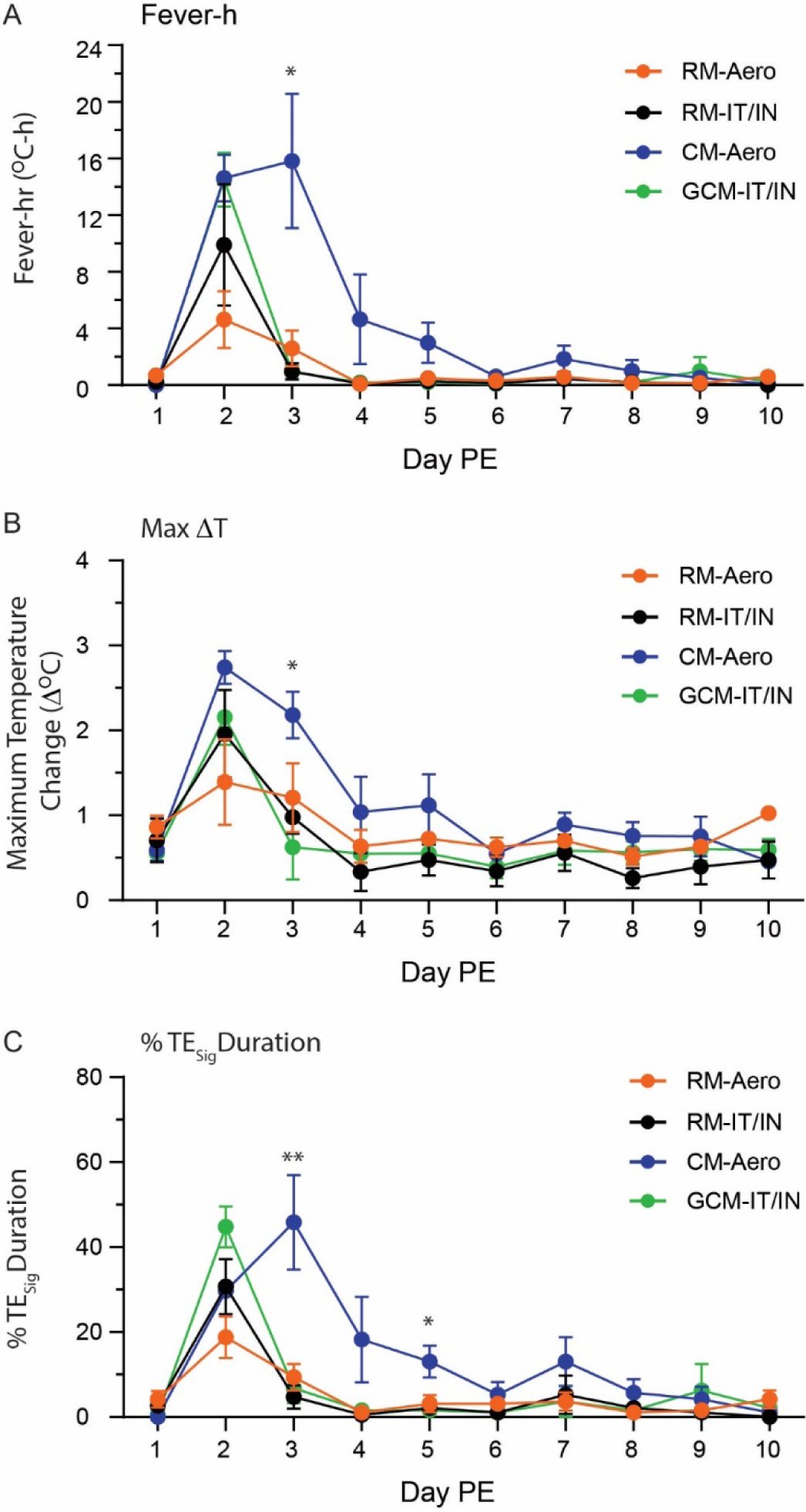
Comparison of significant fever responses during SARS-CoV-2 infection of NHPs. A) Fever-hr B) Maximum daily temperature elevation C) Daily percent TEsig i.e. percent of the 24-hr daily time period when body temperatures were significantly elevated. All data are shown as the group mean ± SEM. A non-parametric one-way ANOVA (Kruskal-Wallis test) was performed for each day.

### Clinical signs of disease following SARS-CoV-2 exposure

SARS-CoV-2 exposure did not result in lethal infection, regardless of species or exposure route. While clinical signs of disease were largely absent upon cage-side examination, evidence of respiratory disease was present on radiographs beginning two days post-exposure (PE) for all animals. At that time, lung infiltrates were limited to animals who were exposed to SARS-CoV-2 via the IT/IN route (2 of 4 for CM IT/IN; 3 of 4 for RM IT/IN). For all groups, the severity of lung disease peaked on Days 4-6 PE, with increases in opacity and worsening of infiltrates (**Figure 3A**). Stabilization and/or mild improvement of the lungs was noted beginning on Day 6 or 8 PE and continuing to the time of euthanasia (Day 9/10 PE), with near complete to complete resolution in all but one animal (CM AE 4). In general, the RM AE animals had the least severe disease, as two of four animals did not have remarkable changes in the lungs during the course of the study (**Figure 3B**). The remaining two RM AE animals had only mild increases in opacity that completely resolved by the end of the study and had radiographic scores that never exceeded a value of two on any given day. Conversely, the CM AE animals developed the most pronounced radiographic changes, despite the fact that the AE animals received a lower dose than the IT/IN animals. Lung consolidation was present in three of four CM AE animals over at least two study days and incomplete resolution of disease by Days 9/10 PE in all animals. A significant difference was observed in radiographic scores for CM AE v RM AE animals on Days 4, 6, and 8 PE, as well as between RM AE v RM IT/IN and between CM AE v CM IT/IN on Day 6 PE (**Figure 3B**).

**Figure 3.**
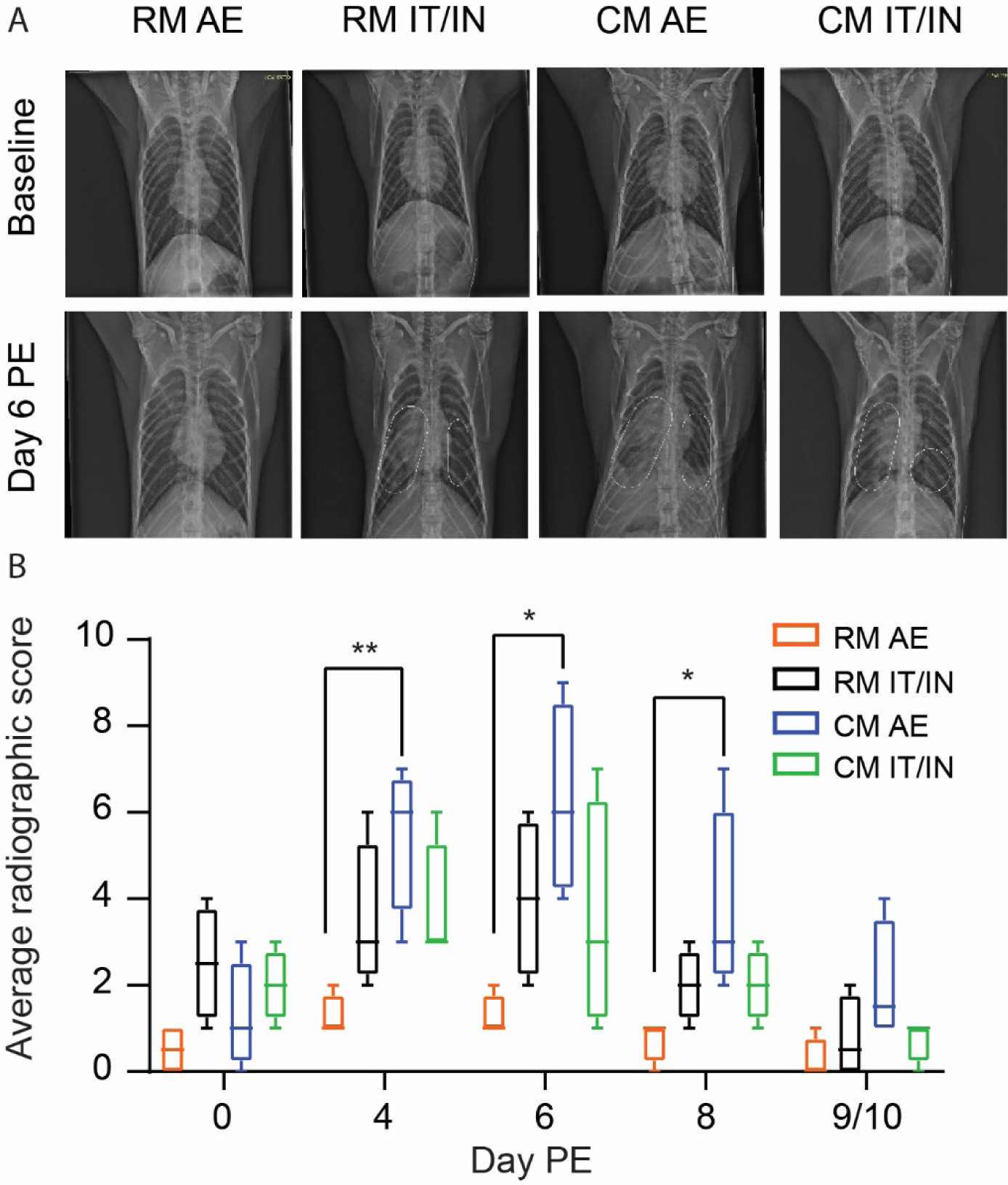
Evidence of clinical disease in radiographs from SARS-CoV-2-infected NHPs. A. Representative radiographs are shown from each group. All images are ventrodorsal. The white dashed lines shown in the Day 6 PE images from RM IT/IN, CM AE, and CM IT/IN outline infiltrates and opacity present in those images; no dashed lines are shown in RM AE due to the absence of lesions. Baseline radiographs were obtained prior to challenge (Day -2). B. Average radiographic scores per group over the course of the study. Box-and-whisker plot showing the range of the data. A non-parametric one-way ANOVA (Kruskal-Wallis test) was performed for each day.

Clinical findings indicative of disease apart from the radiographic findings were largely absent in these animals. However, one consistent finding for RM was erythema of the eyes, which was present between Days 4 and 10 PE (4 of 4 RM IT/IN; 2 of 4 RM AE). This observation was not present in the CMs and the implications of this finding are unclear at this time.

### Clinical Pathology and Immunological Responses

Human COVID-19 disease has been marked by alterations in hematology and serum chemistry markers, including lymphopenia and increases in C-reactive protein (CRP) (3–6). Other markers, such as creatine kinase (CK), aspartate aminotransferase (AST) and alanine aminotransferase (ALT), have been associated with more severe cases of disease and poor prognosis for survival (19–20). While there was a great degree of animal-animal variation in clinical pathology responses that limited statistical significance, a number of trends did emerge. A decrease in platelet counts was observed early (Day 2 PE) following SARS-CoV-2 exposure, with partial recovery by the day of terminal sampling (Day 9/10 PE) (**Figure 4A**). Most groups demonstrated a decrease in lymphocyte counts on Day 2 PE, which was followed by a gradual increase to baseline or above baseline over the remainder of the in-life study period (**Suppl Figure 2A**). One of the most dramatic hematological changes observed was monocytosis, with peak changes in excess of +3100% between Days 6 and 9/10 PE (**Suppl Figure 2B**).

**Figure 4:**
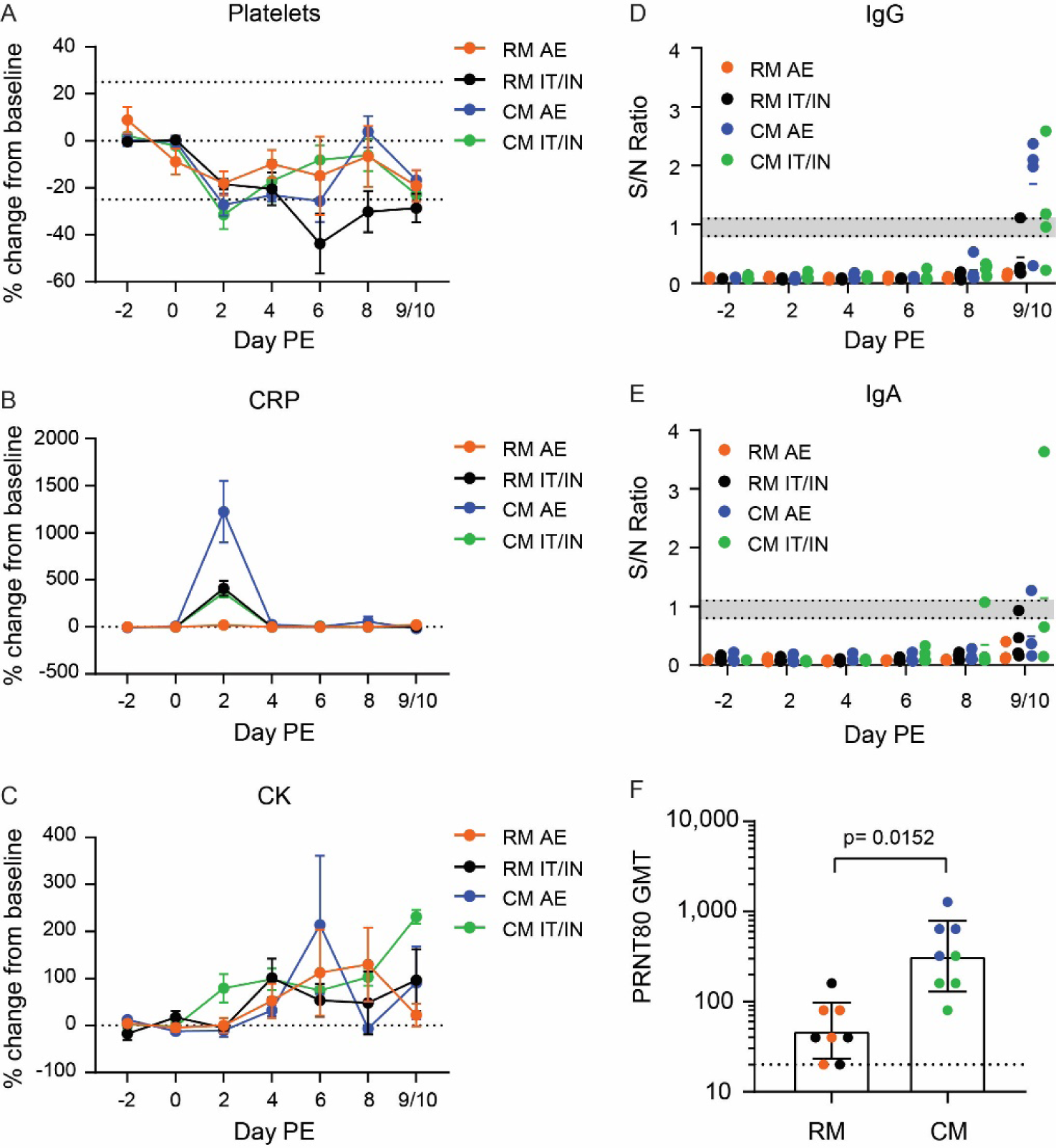
Clinical pathology and serological responses to SARS-CoV-2 exposure. A-C. Percent change from baseline for A) platelets, B) CRP, and C) CK over the course of the study. The dashed line at y=0 represents the baseline, which is defined as the average of the results from Days -2 and 0. Values greater than or equal to a 25% change from baseline (dashed lines in A) were considered noteworthy. Data are shown as mean ± SEM. D-E. IgG (D) and IgA (E) results from the Euroimmun SARS-CoV-2 ELISA kits. Results are shown as the signal to noise ratio. The bottom and top dotted lines represent the assay cutoffs for negative and positive results respectively, with the gray shaded region representing indeterminate results. The mean is denoted by a colored dash for each group F) PRNT80 geometric mean titer (GMT) for RM versus CM in serum obtained at the terminal time point (Day 9/10 PE), with each animal represented as a dot and color corresponding to experimental group. The dotted line represents the assay cutoff for positive results (PRNT80 of 20). Error bars represent the geometric SD. Unpaired t test was performed, with * p<0.05.

With the exception of the RM AE group, increases in one or more hepatocellular enzymes (ALT and AST) were routinely observed for animals in the RM IT/IN, CM AE, and CM IT/IN groups starting as early as two days PE; the RM AE animals had very mild changes in these enzymes (**Suppl Figures 2C and 2D**). The most noteworthy and consistent serum chemistry alterations were highly elevated levels of C-reactive protein (CRP) and creatine kinase (CK) post-exposure to SARS-CoV-2 (**Figures 4B and 4C**). CRP levels increased in nearly all NHPs in the RM IT/IN, CM AE, and CM IT/IN groups beginning two days PE. Only 2/4 RM AE animals demonstrated increases in CRP (+36-46%), which were mild and significantly lower than the other groups (+190-1786%). In the IT/IN animals (RM and CM), CRP levels were moderately elevated and largely returned to baseline by the next sampling time point (Day 4 PE). The highest levels of CRP were observed in the CM AE group, with >1000-fold increases in 3/4 CM AE animals on Day 2 PE. As seen with the IT/IN animals, CRP levels returned to baseline by Day 4 PE in three of the four CM AE animals. Similar elevations were observed in CK levels, although these alterations persisted for a longer duration (often until the day of euthanasia) and included the RM AE group as well.

Serum was also evaluated for antibody responses to SARS-CoV-2 infection using the commercially available Euroimmun ELISA kit for IgG and IgA. An IgG response was only detectable at the terminal time point (Days 9 or 10 PE) in select animals from the CM AE and CM IT/IN groups (**Figure 4D**). This finding was not unexpected as a previous study noted detectable IgG responses by Day 10 PE in CMs and Day 15 PE in RMs (13). Similarly, an IgA response was only detected in two CMs by Day 9/10 PE (**Figure 4E**).

A MAGPIX multiplex assay for IgG and IgM responses directed against SARS-CoV-2 full-length spike glycoprotein, receptor-binding domain (RBD) of the spike protein, and nucleoprotein provided additional insight into the antibody response following SARS-CoV-2 exposure (**Suppl Figure 3**). IgM responses to SARS-CoV-2 glycoproteins were detected as early as Day 6 PE, while IgG responses were detected by Day 8 PE. As previously observed (13), the IgG assay, particularly in the CMs, had less specificity for the glycoproteins than IgM, as nucleoprotein responses were detected in a subset of animals. IgM responses predominantly favored the full spike protein over the RBD, with responses appearing earlier in CMs than RMs. IgG responses were more evenly distributed between the full spike protein and RBD, with the highest responses favoring the IT/IN route over AE.

Terminal sera samples (Day 9 or 10 PE) were assessed in a plaque reduction neutralization test (PRNT) for neutralizing antibodies directed against SARS-CoV-2. An obvious difference in neutralizing antibody titers between the two species emerged, as all but one RM had PRNT80 titers of 80 or lower (**Figure 4F**). The majority of the CMs demonstrated stronger neutralizing antibody responses, with PRNT80 values ranging from 160 to 1280. The PRNT80 titers were significantly greater in CMs versus RMs, with the three highest titers attributed to the CM AE group.

### Pathology

All NHPs were euthanized on either Day 9 or Day 10 PE for histopathologic analysis. These days were selected with the intention of viewing SARS-CoV-2-induced respiratory pathology prior to complete resolution. No significant gross findings were noted except that one animal from the CM AE group had pulmonary fibrinous adhesions from the thoracic cavity to the pleural surface of the lungs (**Figure 5A**). Moreover, more than half of the animals from each group had enlarged tracheobronchial lymph nodes of varying degrees of severity.

**Figure 5.**
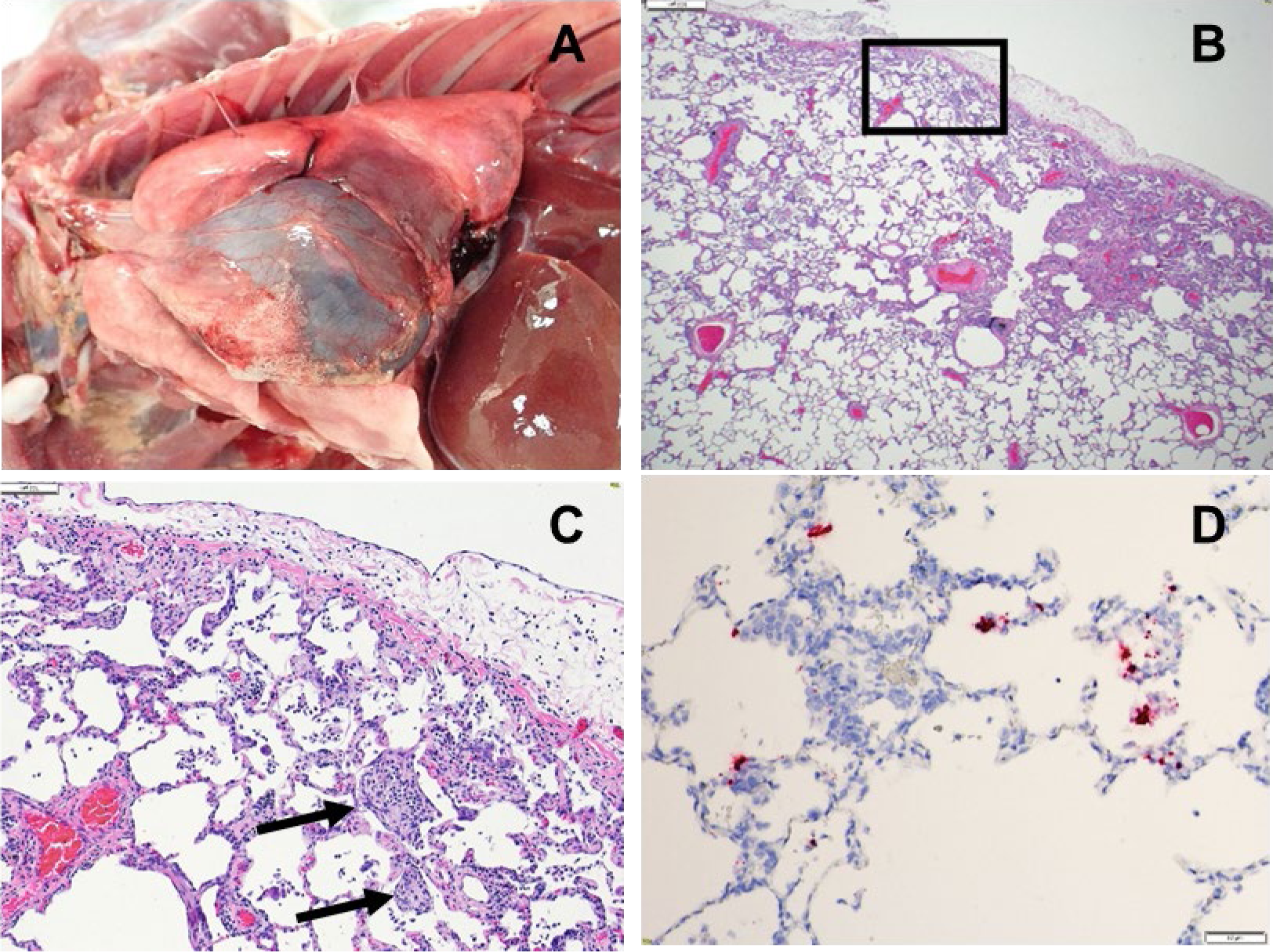
Respiratory pathology in CM exposed to SARS-CoV-2 by the AE route. Images shown from animal CM AE 1. A. Thoracic cavity: Multiple fibrinous adhesions between left cranial and caudal lung lobes and the thoracic wall. B. Lung, left caudal lung lobe, peripheral: Multifocal, moderate, interstitial pneumonia with pleural fibrin, 2x, H&E. C. Lung, higher magnification of boxed area in B: Multifocal moderate lymphohistiocytic interstitial pneumonia with type II pneumocyte hyperplasia, intra-alveolar fibrin deposition (black arrow), septal fibrosis, pleuritis and pleural fibrin, 10x, H&E. D. Lung: ISH positive in areas of inflammation, 20x, RNA probe for SARS-CoV-2.

Histological findings are present in the lungs of all groups that are compatible with lesions that have been described in humans infected with SARS-CoV-2 (7). Significant pulmonary histological lesions for individual animals in the four groups are detailed in Suppl. Table 1 and include inflammation, type 2 pneumocyte hyperplasia, multinucleated giant cells, alveolar fibrin deposition (septal and intra-alveolar) and septal fibrous change (**Figure 5B–C, Suppl. Figures 4-6**). The lungs of the CMs were more severely affected compared to RMs (**Suppl. Table 1**).

**Figure 6.**
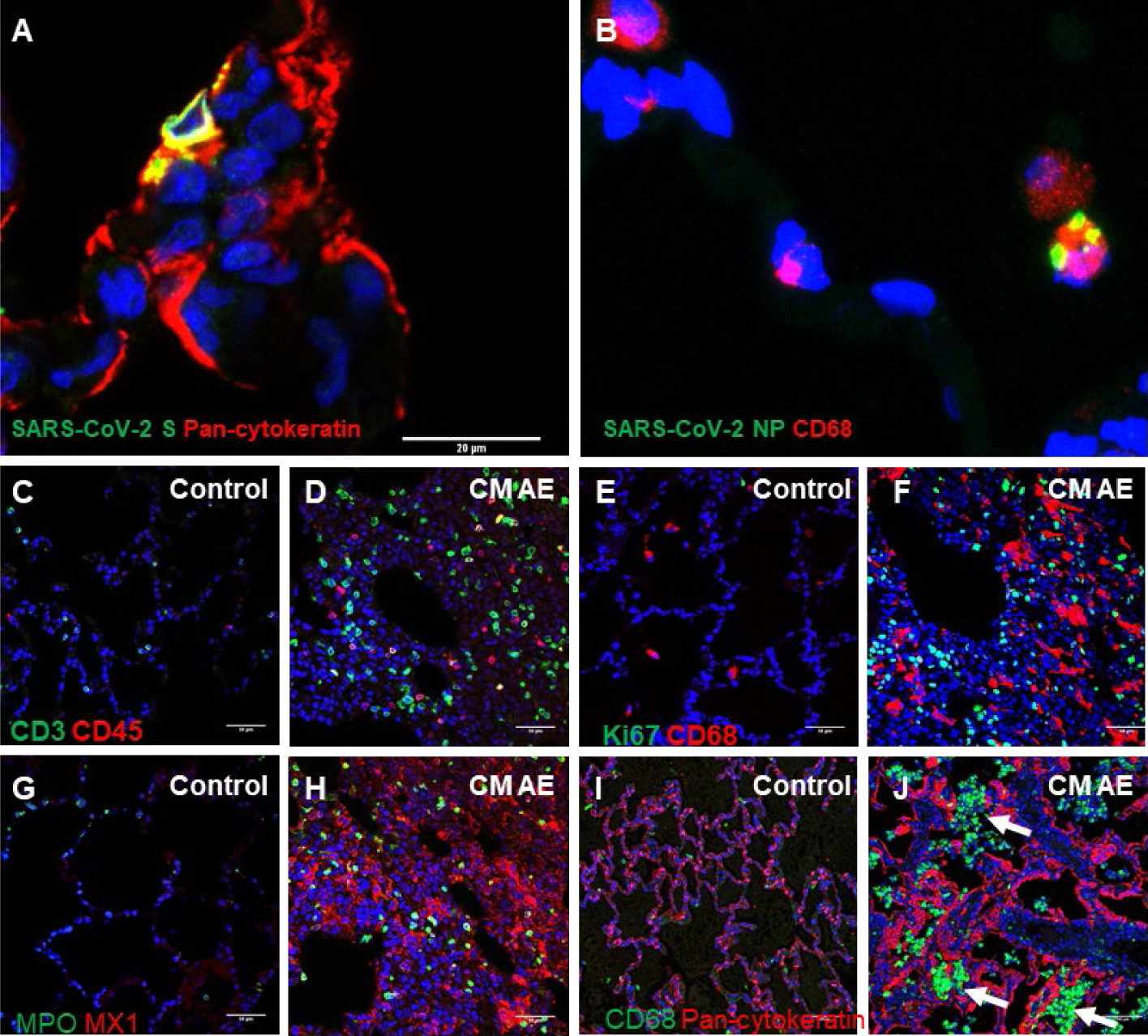
The cellular targets of SARS-CoV-2 and inflammatory infiltrates in the lungs of CM AE NHPs. A. The spike (S) protein of SARS-CoV-2 (green) was detected in pneumocytes (red) labelled by anti-Pan-cytokeratin antibody (red). B. The nucleoprotein (NP) of SARS-CoV-2 (green) was detected in CD68+ macrophages (red). C–H. In comparison to uninfected control lung tissue, CD3+ T cells (green in C and D), CD45+ leukocytes (red in C and D), CD68+ macrophages (red in E and F), and Ki67+ proliferating cells (green in E and F), MPO+ polymorphonuclear cells (neutrophils, eosinophils, and basophils, green in G and H), expression of the type 1 interferon-induced GTP-binding protein Mx1 (a readout of type 1 interferon response, red in G and H) in the lungs of CM AE NHPs exposed to SARS-CoV-2. I–J. Alveolar spaces were congested by CD68+ macrophages (green, arrows) in some areas of the lungs of CM AE animals. Lung epithelium is stained outlined by anti-Pan-cytokeratin antibody (red). Nucleus was stained by DAPI (blue).

Additionally, the severity of pulmonary lesions were greatest in the CM AE group. These lesions were confirmed to be the result of SARS-CoV-2 infection using RNA *in situ* hybridization (ISH) to detect genomic RNA of SARS-CoV-2 (**Figure 5D**). The enlarged tracheobronchial lymph nodes noted during gross examination corresponded to lymph node hyperplasia. This finding, along with nasal turbinate inflammation and edema, are suspected to be disease-related/exposure- related.

Since CM AE animals had the most severe pulmonary pathology as compared to the other groups, we performed immunofluorescence staining to map the cellular targets of SARS-CoV-2 and characterize the inflammatory infiltrates in the lungs of CM AE animals.

Immunofluorescence staining demonstrated SARS-CoV-2 primarily infected pan-Cytokeratin- labelled pneumocytes and CD68+ macrophages (**Figure 6A-B**). Consistent with the above histopathological observations, increased number of CD3+ T cells, CD45+ leukocytes, CD68+ macrophages, Ki67+ proliferating cells, and myeloperoxidase (MPO)+ polymorphonuclear cells (neutrophils, eosinophils, and basophils) were present in the lung of CM AE animals in comparison with the lung of historic uninfected CMs (**Figure 6C-H**). Furthermore, significantly increased expression of the type 1 interferon-induced GTP-binding protein Mx1, was detected in the lungs of CM AE animals (**Figure 6G-H**). Interestingly, immunofluorescence staining illustrated alveolar spaces were congested by CD68+ macrophages in some areas of the lung of cynomolgus macaques with aerosol exposure of SARS-CoV-2 (**Figure 6I-J**). Together, the above pathologic data indicates CM AE animals had the most consistent and severe pulmonary lesions among the four groups of nonhuman primates.

## Discussion

The potential for a previously unknown infectious agent to rapidly encircle the globe became reality at the close of 2019, with the identification of COVID-19 and its causative agent SARS-CoV-2. The rapid spread from the outbreak in China into a global pandemic spurred the development of the necessary tools to fight the pandemic, including relevant animal models that reproduce the hallmarks of human disease. While the highest aspiration of animal model development is a single model that is identical to the human condition, the reality is typically a series of models, each capturing a different combination of key disease features and/or severity. To date, the nonhuman primate models for SARS-CoV-2 have primarily replicated the mild form of COVID-19 that predominates in humans. However, with a few exceptions, these models have yet to capture the more severe end of the clinical spectrum of human COVID-19 disease.

Previous NHP studies with SARS-CoV-2 have explored differences in primate species as well as in routes of virus exposure (10, 13). Here, we expanded on this work by performing a direct comparison of two routes of exposure (AE and combined IT/IN) in two NHP species with the goal of refining the NHP model for SARS-CoV-2 and recapitulating some of the more significant clinical and pathological effects of SARS-CoV-2 infection.

Exposure of RM and CM to SARS-CoV-2 via either AE or combined IT/IN administration resulted in mild disease that was similar to human COVID-19. Both routes of exposure enabled delivery of virus to the lower and upper respiratory tract using two different processes: the IT/IN route used direct contact of the nasal and respiratory tissues with small volumes of virus-containing liquid, while the AE route utilized inhalation of experimentally-generated aerosols containing virus. Although the aerosolization process inherently results in an approximate two log decrease in exposure dose, inhalational exposure to aerosols is likely more physiologically relevant than direct IT instillation (21). Additionally, the small size of the aerosol particles produced (1-3 µm) generates a distinct distribution pattern from liquid or large droplets, as small particles are capable of traveling into deep lung tissue (18, 22).

All CMs on the study displayed evidence of either temperature elevation or fever by telemetry, although the two routes of exposure produced different patterns. Fever in the CM IT/IN animals was short in duration, while the CM AE animals demonstrated fever that often persisted for several days. Fever was present in select RM animals as well, although this clinical finding was less consistent and lower in intensity. While a previous comparison of CMs and RMs exposed to SARS-CoV-2 did not note any significant temperature changes (10), another study also utilizing real-time telemetry was able to detect different temperature responses amongst the various NHP species (13); this highlights the value of using such technology to monitor physiological parameters that might be otherwise overlooked on physical examination. The consistent presentation of fever in the CMs, particularly the intensity and duration of fever in AE animals, presents an attractive endpoint for this model, particularly in light of the absence of mortality. In non-lethal models, alternative endpoints such as fever and clinical pathology become even more important in assessing the efficacy of vaccines or therapeutics. As such, protective effects may be more readily apparent in CMs (than RMs) exposed to SARS-CoV-2, and even more so when aerosol exposure is used instead of IT/IN. Moreover, the stronger immunological response in CMs over RMs, as measured by Magpix, ELISA, and PRNT, following SARS-CoV-2 infection may have additional implications for vaccine studies where these measurements may be potential correlates of protection.

The decision to euthanize the animals on Days 9 and 10 PE enabled the evaluation of the lungs prior to complete resolution of disease and the resulting pathology. While the ISH results from a previous SARS-CoV-2 NHP study were all negative at the time of necropsy on Study Day 18 (13), the majority of CM and a few RM were still ISH positive in the lung tissue on Days 9 and 10 PE in this study. This suggests that earlier scheduled termination may be useful in assessing the protective vaccine or therapeutic effect on lung pathology in these animals. The CM AE animals displayed the most severe pathology of all the groups at the time of necropsy, with moderate levels of inflammation, type II pneumocyte hyperplasia, fibrin deposition, and fibrosis. By comparison, inflammation in the RM (both AE and IT/IN) was minimal, with little to no fibrosis present. While Salguero *et al* found similar degrees of respiratory pathology in both cynomolgus and rhesus macaques exposed to SARS-CoV-2 by IT/IN (10), the addition of AE exposure here appears to result in very different presentations in the two species. While the ability of small aerosol particles to reach deeper sections of the lung may be an explanation of the severity observed in the CM AE animals, it is interesting that the same degree of severity is not present in the RM AE group. It is possible that species-specific differences in immune cell composition and/or tissue structures may explain this dichotomy. Although we hypothesize that the increased severity of pathology in the CM AE animals may be attributable to particle size and penetration, it is difficult to make any definitive conclusions due to the lack of longitudinal pathology samples in this study, as the respiratory pathology was already well-established at Days 9/10 PE. A serial sampling study in which the progression of SARS-CoV-2 infection through the respiratory tract is monitored over time would provide valuable information regarding the influence of exposure route (AE v IT/IN) on the resulting distribution of pathology in the upper versus lower respiratory tract.

Similar to the previous USAMRIID SARS-CoV-2 study conducted in NHPs (13), erythema around the eyes was a consistent finding in this study and was only present in the RM. As small particle aerosol exposure utilizes a head-only chamber in which the eyes are not covered during aerosol exposure, it was previously hypothesized that ocular exposure to SARS-CoV-2 during the aerosolization process may have been responsible for this observation in the previous study (13). However, the fact that this clinical sign was present in all of the RM IT/IN animals and only half of the RM AE animals in this study, suggests that another mechanism may be responsible and should be investigated further. Moreover, this clinical observation was not noted in AGMs exposed to SARS-CoV-2 by AE (11).

Although the CM AE animals developed the most severe pathology and physiological changes, exposure of both RM and CM to SARS-CoV-2 via either aerosol or combined IT/IN administration resulted in mild disease that was similar to human COVID-19 (**Figure 7**). One of the unexpected effects of the SARS-CoV-2 pandemic has been an unprecedented demand for nonhuman primates for COVID research. This increased demand, coupled with decreased supply due to export restrictions, is quickly resulting in a shortage of available NHPs.

**Figure 7.**
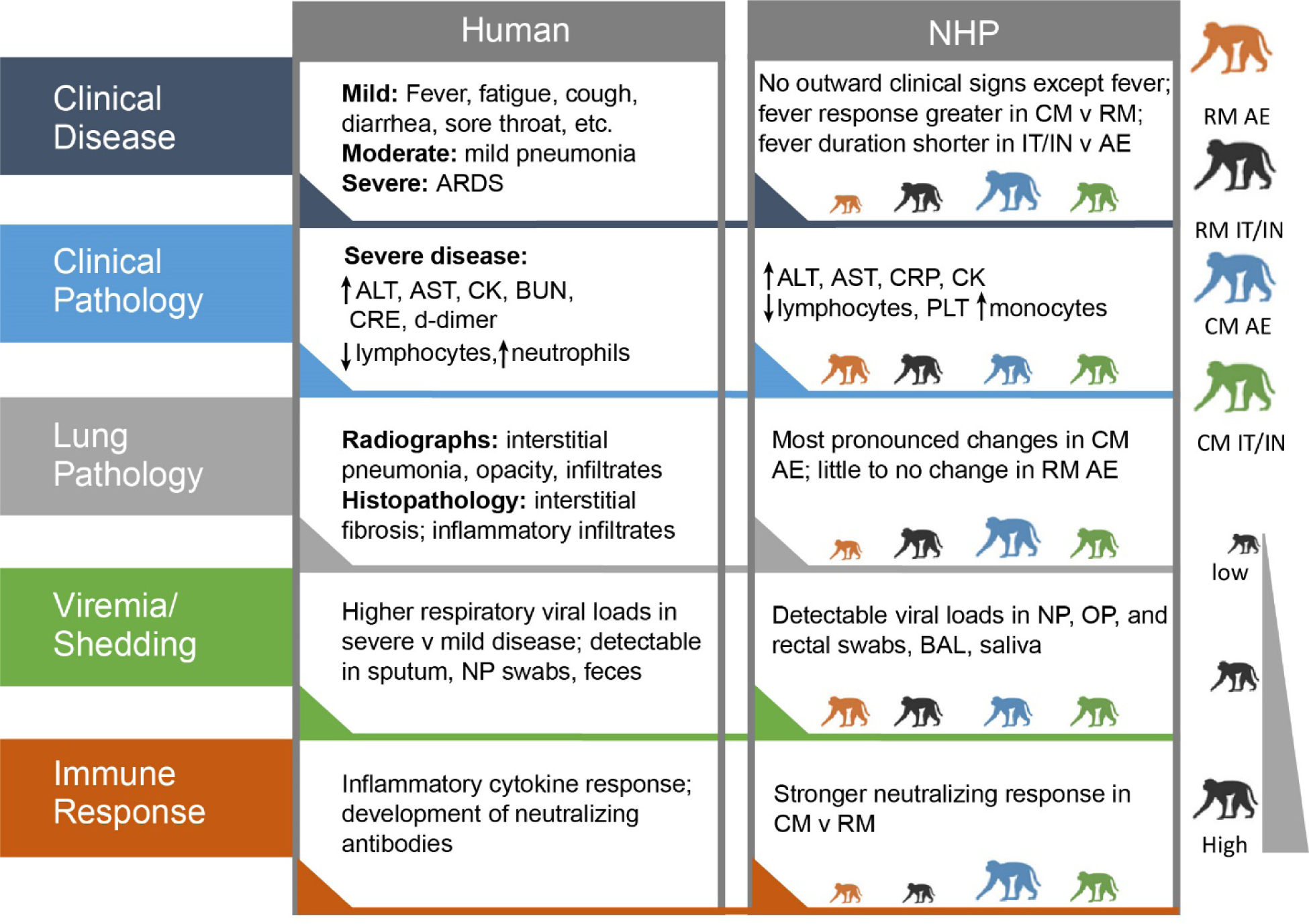
Comparison of human COVID-19 and SARS-CoV-2 disease in NHPs. The size of the NHPs shows the relative response for that group.

Additionally, aerosol exposure capability requires specialized equipment and expertise that is not available at all research institutions. The IT/IN route is an alternative that presents fewer technical challenges and can more readily be implemented at a wide range of laboratories. This study demonstrates that different NHP species and routes of exposure produce a spectrum of disease severity, which may be comparable to the range of COVID-19 disease presentations in the human population, and moreover, affords the SARS-CoV-2 research community several NHP model options that can be tailored to facility capabilities and animal availability.

Taken together, the clinical findings suggest that CM may be the better model for SARS-CoV-2 than RM, as these animals had the most severe and consistent disease presentation. Of the two exposure routes, AE produced more severe respiratory disease and lung pathology than the IT/IN route in the CM. While the AE animals received an exposure dose that was approximately two logs lower than that of the IT/IN animals, the ability of small particle aerosol to reach deep into the lung tissue may be a possible explanation for this finding. It is interesting to note that while the AE animals received a lower dose of virus, SARS-CoV-2 was detected at similar levels in all four groups overall, as indicated by PCR (N2 and subgenomic) and plaque assay. If anything, the CM AE animals, which had the most severe pathology, demonstrated a trend toward lower levels of infectious and replicating virus than the other groups, although this did not achieve statistical significance due to animal variability. Future studies that are able to expose animals to a higher AE dose of SARS-CoV-2 may result in further refinement of this promising NHP model, potentially with a concomitant increase in disease severity and/or mortality.

## Materials and Methods

### Virus

A stock of SARS-CoV-2, Washington state first isolate in 2020 (WA-1/2020), was obtained from the Centers for Disease Control and Prevention and designated as Lot R4713. This strain was isolated from nasopharyngeal and oropharyngeal swabs obtained from a patient in Washington state who had traveled to China (23). The CDC isolate had been passaged three times in CCL-81 cells prior to receipt at the USAMRIID lab. A master (Lot R4714a) and seed (Lot R4716) were made. The seed stock was then passaged in ATCC Vero 76 cells at an MOI of 0.01 and incubated for approximately 50 hours. The supernatant was clarified by centrifugation and the resulting virus production stock was designated as Lot R4719. The production stock, contained an average of 5.45×10^6^ pfu/mL of infectious virus as determined using a neutral red plaque assay. R4719 was fully sequenced using an Illumina MiSeq platform. The production stock underwent additional testing to evaluate sterility, mycoplasma and endotoxin levels, as well as a number of real-time reverse transcriptase polymerase chain reaction (RT-PCR) assays for exclusivity and inclusivity, to include two specific for SARS-CoV-2 virus. Lot R4719 was determined to have no detectable mycoplasma, endotoxin or adventitious agents based on the assays and techniques used. No known contaminants were detected during sequencing of the stock. Identity was confirmed by real-time RT-PCR and sequencing; the sequence of lot R4719 was identical to the original stock obtained from the patient isolate, with no deletion or mutations observed.

### Telemetry

Telemetry implants were utilized for continuous monitoring of body temperature and activity in the NHPs. Approximately ten days prior to virus exposure, animals were surgically implanted with M00 implants (Data Sciences International, Inc., St. Paul, MN) by USAMRIID veterinary staff and allowed to recover from surgery prior to release onto study. Animals with implanted devices were housed in standard NHP caging. The implants multiplexed raw data signals from the sensors and wirelessly transmitted digital signals at a preset frequency to the receivers. The digital signals were received by multiple TRX receivers, then transmitted over Cat 5e cable to communication link controllers (CLC). The signals were then routed over Cat 5e cable to data acquisition computers, which captured, reduced and stored the digital data in data files (i.e., NSS files) using the Notocord-hem Evolution software platform (Version 4.3.0.77, Notocord Inc., Newark, New Jersey). Temperature and activity signals were collected using a sampling rate of one sample per second.

### NHPs

The study used eight healthy cynomolgus macaques and eight healthy rhesus macaques with equal distribution of sex among the groups (n=8 males and n=8 females; n=2 of each per group). All animals were of Chinese origin and were obtained from the USAMRIID NHP colony. The animals were between the ages of 5.3-9.8 years and weighed between 2.976 and 9.585 kg. Animals were determined to be serologically negative for SARS-CoV-2 at the outset of the study as determined by ELISA and PRNT (described below).

### Virus exposure

Animals were randomized to exposure route balanced by sex and weight. On the designated day of virus exposure, animals were exposed to the WA-1/2020 strain of SARS-CoV- 2 by either the small particle AE route (n=8) or combined IT/IN administration (n=8).

The AE exposure dose for each animal was calculated from the minute volume determined with a plexiglass whole body plethysmograph box using Buxco FinePointe software. The total volume of aerosol inhaled was determined by the exposure time required to deliver the estimated inhaled dose. Animals were exposed to the target AE dose between 5.0×10^4^ and 5.0×10^5^ pfu of challenge agent in the USAMRIID head-only exposure system. The AE challenge was generated using a Collison Nebulizer to produce a highly respirable aerosol (flow rate 7.5±0.1 L/minute). The system generates a target aerosol of 1 to 3 µm mass median aerodynamic diameter determined by aerodynamic particle sizer. Samples of the aerosol collected from the exposure chamber using an all-glass impinger (AGI) during each challenge was assessed using a neutral red plaque assay to determine the inhaled dose for each animal.

The combined IT/IN animals were administered a target dose of 2.0×10^7^ pfu split between the two routes of exposure. Four mL of virus was administered with a syringe via feeding tube catheter for the IT route. Following administration of virus, a bolus of air was delivered using a syringe to ensure that the full dose of virus was administered and that no material remained in the catheter. Immediately following IT administration, animals were administered 0.25 mL of virus inserted dropwise into each nare via syringe for the IN route (a total of 0.5 mL for IN route). The animals’ heads were held facing upwards for up to 2 minutes to ensure proper delivery of the inoculum into the nasal passages. A neutral red plaque assay was performed on the viral inoculum to confirm the titer of the stock virus.

### Post-exposure observations and sample collection

Animals were observed at least once daily for clinical signs of SARS-CoV-2 infection, including respiratory signs and changes in responsiveness and activity. Physical examinations to include weights and radiographs were performed under anesthesia two days prior to virus exposure (Day -2), the day of exposure (designated here as Day 0), and days 2, 4, 6, 8, and at the time of euthanasia (9 or 10) post-exposure (PE). While under anesthesia, blood and nasopharyngeal, oropharyngeal, and rectal swabs were collected for evaluation of viral RNA, infectious virus, and clinical pathology to include serum chemistry and hematology. Swabs were collected into 1 mL of viral transport media (VTM; Hanks Balanced Salt Solution containing 2% heat-inactivated fetal bovine serum, 100 µg/mL gentamicin, and 0.5 µg/mL amphotericin B), vortexed for 15-20 seconds, and the lysate was removed. Eight animals (two from each of the four groups) were randomly selected for euthanasia on Day 9 PE, with the eight remaining animals euthanized on Day 10 PE.

### Radiographs

Ventrodorsal and lateral radiographs were performed during each anesthetized physical examination and were scored for evidence of respiratory disease. Radiographic scores for each animal represent the summation of scores across each lung lobe per day.

|  |  |
| --- | --- |
| 0 | Normal, no change from baseline |
| 1 | Mild opacity increase and/or mild interstitial pulmonary infiltrates |
| 2 | Moderate pulmonary infiltrates +/- small areas of consolidation |
| 3 | Severe pulmonary infiltrates, large areas of consolidation, alveolar pattern |

### Clinical pathology

Serum chemistry analysis was performed on a Piccolo point-of-care instrument (Abaxis) using the General Chemistry 13 and MetLyte Plus CRP panels. Hematology analysis was performed on EDTA whole blood using a VETSCAN® HM5 hematology analyzer and multispecies software.

### SARS-CoV-2 quantitative real-time RT-PCR

Samples collected for PCR analysis were inactivated in TRIzol™ LS in a ratio of 3 parts TRIzol™ LS to one part sample. Inactivated samples were then extracted and eluted with AVE buffer using a QIAamp® Viral RNA Mini Kit IAW SOP AP-09-31. The RT-PCR reaction used Invitrogen™ SuperScript® One-Step RT-PCR System with additional magnesium sulfate (MgSO4) added to a final concentration of 3.0 mM. Specimens were run in triplicate using a 5 µL volume on an Applied Biosystems® 7500 Fast Dx instrument. The average of the triplicates were multiplied by 200 to obtain genomic equivalents per mL, then multiplied by a dilution factor of 4 (1 part plasma to 3 parts TRIzol™ LS) for the final reported value. The genomic equivalents were determined using a standard curve of synthetic RNA of known concentration. The sequences of primers and probes for the N2 gene from SARS-CoV-2 that were used in the assay were previously described in (13).

### SARS-CoV-2 subgenomic RNA real-time RT-PCR

Total SARS-CoV-2 E gene and subgenomic E gene target copy numbers were determined by real-time RT-PCR using previously described assays (24–25) and a synthetic RNA containing the subgenomic E RNA sequence (Bio-Synthesis, Inc., Lewisville, TX, USA). Extracted nucleic acid was tested in triplicate (5 μL extracted nucleic acid) with each assay using the synthetic RNA as a standard curve on each run. Samples were run on the LightCycler 480 (Roche Diagnostics, Indianapolis, IN, USA) using Superscript III one-step RT-PCR system with Platinum taq (Thermo Fisher Scientific). Cycling conditions were: 50°C for 10 min; 95°C for 3 min; 45 cycles of 95°C for 10 sec, 56°C for 15 sec, and 72°C for 5 sec; and a final hold of 40°C for 30 sec. Copy numbers for each target were determined using the synthetic RNA standard curve, and the amount of target amplicon in the original sample was calculated from these results. The limit of detection for the assay is 100 copies/µL.

### Plaque assay

A neutral red plaque assay using ATCC Vero 76 cells was performed on the virus stock, AGIs from the AE-exposed animals, and nasopharyngeal and oropharyngeal swab lysates as previously described (13). Briefly, the required dilutions of each specimen, in duplicate, were added to plates containing ATCC Vero 76 cells on assay day 0. The cells were stained with neutral red on assay day 2, and plaque counts were obtained on assay day 3.

### Euroimmun SARS-CoV-2 S1 ELISA

NHP serum samples were screened with the Euroimmun SARS-CoV-2 S1 ELISA (Euroimmun, EI 2606-9601 G) as per kit instructions. Briefly, the kit materials were brought to room temperature for 30 minutes. Serum samples were diluted 1:101 using the supplied sample buffer. 100 µL of the diluted samples, supplied controls, and supplied calibrator were added to the pre-coated wells and incubated at 37°C for 1 hour. After 1 hour, the plate was washed 3 times with 300 uL of supplied wash buffer using a microplate washer (Biotek 405TS). 100 µL of enzyme conjugate was added to the wells and incubated at 37°C for 30 minutes. The plate was washed 3 times as outlined above prior to adding 100 µL of substrate for 30 minutes at RT. Finally, 100 µL of stop solution was added prior to reading absorbance at 450 nm, with a reference wavelength at 635 nm (Tecan M200). Data was processed according to kit instructions to determine negative, positive, or borderline results.

### SARS-CoV-2 MAGPIX multiplex immunoassay

Serum samples were diluted at 1:100 phosphate buffer saline (PBS) with 0.02% Tween- 20 (PBST) with 5% skim milk (PBST-SK). Recombinant SARS-CoV-2 full trimeric spike (gift from Dr. Jason McLellan’s group; UT-Austin) (26), RBD (Sino Biological, 40592-V08H), and NP (Native Antigen Company, REC31812-100) proteins were coupled to Magplex microsphere regions #45, #65, and #25 using the Luminex xMAP® antibody coupling kit (Luminex Inc., Austin, TX, USA) according to the manufacturer’s instructions. Each individual antigen-coupled bead was mixed at a 1:1 ratio prior to diluting in PBST to 5 × 10^4^ microspheres/mL and added to the wells of a Costar polystyrene 96-well plate at 50 µL per well (2500 microspheres of each antigen bead set/well). The plate was placed on a magnetic plate separator (Luminex Inc.) covered with foil, and microspheres were allowed to collect for 60 sec. While still attached to the magnet, the buffer was removed from the plate by inverting and disposing into the sink. Then, 50 µL of the diluted plasma samples were added to appropriate wells. The plate was covered with a black, vinyl plate cover and incubated with shaking for 1 hr at RT. The plate was washed three times with 100 µL of PBST for each wash, using the plate magnet to retain the Magplex microspheres in the wells. Liquid was discarded as above. 50 µL of a 1:100 dilution of mouse anti-human IgM phycoerythrin conjugate (Invitrogen, MA1-10381) or goat anti-human IgG phycoerythrin conjugate (Sigma, P9170) in PBST-SK was added to the wells. The plate was covered again with a black, vinyl plate sealer and incubated with shaking for 1 hr at RT. After incubation, the plate was washed three times as detailed above and the Magplex microspheres were resuspended in 100 µL of PBST for analysis on the Magpix instrument. Raw data was reported as median fluorescence intensity for each bead set in the multiplex.

### Plaque reduction neutralization test (PRNT)

The PRNT was performed on serum samples from the terminal time point as previously described (13). PRNT80 titers were calculated as the reciprocal of the highest dilution that generated an 80% reduction in plaque counts relative to virus only.

### Necropsy and histology

Necropsies were conducted by a veterinary pathologist on all animals in this study. The tissue samples were trimmed, routinely processed, and embedded in paraffin. Sections of the paraffin-embedded tissues 5 um thick were cut for histology. For histology, slides were deparaffined, stained with hematoxylin and eosin (H&E), coverslipped, and labeled.

### Immunofluorescence

Formalin-fixed paraffin embedded (FFPE) tissue sections were deparaffinized using xylene and a series of ethanol washes. After 0.1% Sudan black B (Sigma) treatment to eliminate the autofluorescence background, the sections were heated in Tris-EDTA buffer (10 mM Tris Base, 1 mM EDTA Solution, 0.05% Tween 20, pH 9.0) for 15 minutes to reverse formaldehyde crosslinks. After rinses with PBS (pH 7.4), the section were blocked with PBT (PBS +0.1% Tween-20) containing 5% normal goat serum overnight at 4°C. Then the sections were incubated with primary antibodies: rabbit anti-SARS-CoV Spike (1:200, 40150-T62-COV2, Sino Biological, Chesterbrook, PA, USA), mouse anti-SARS-CoV NP (1:200, 40143-MM05, Sino Biological), mouse anti-Pan-Cytokeratin (1:100, M351529-2, Dako Agilent Pathology Solutions), rabbit anti-CD3 (1:200, A045229-2, Dako Agilent Pathology Solutions, Carpinteria, CA, USA), rabbit anti-MPO (1:200, A039829-2, Dako Agilent Pathology Solutions, Carpinteria, CA, USA), rabbit anti-CD68 (1:200, ab125047, Abcam, Cambridge, MA, USA), mouse anti-CD68 (1:100, M081401-2, Dako Agilent Pathology Solutions), mouse anti-CD45 (1:100, M070101-2, Dako Agilent Pathology Solutions), and/or mouse anti-MX1 (1:200, MABF938, Millipore Sigma, Burlington, MA, USA) for 2 hours at room temperature. After rinses with PBT, the sections were incubated with secondary goat anti-rabbit Alexa Fluor 488 (1:500, Thermo Fisher Scientific) and goat anti-mouse Alexa Fluor 568 (red, 1:500, Thermo Fisher Scientific) antibodies, for 1 hour at room temperature. Sections were cover slipped using the Vectashield mounting medium with DAPI (Vector Laboratories). Images were captured on a Zeiss LSM 880 confocal system (Zeiss, Oberkochen, Germany) and processed using ImageJ software (National Institutes of Health, Bethesda, MD).

### RNA *in situ* hybridization

To detect SARS-CoV-2 genomic RNA in FFPE tissues, in situ hybridization (ISH) was performed using the RNAscope 2.5 HD RED kit (Advanced Cell Diagnostics, Newark, CA, USA) as described previously (27). Briefly, forty ZZ ISH probes targeting SARS-CoV-2 genomic RNA fragment 21571-25392 (GenBank #LC528233.1) were designed and synthesized by Advanced Cell Diagnostics (#854841). Tissue sections were deparaffinized with xylene, underwent a series of ethanol washes and peroxidase blocking, and were then heated in kit-provided antigen retrieval buffer and digested by kit-provided proteinase. Sections were exposed to ISH target probe pairs and incubated at 40°C in a hybridization oven for 2 h. After rinsing, ISH signal was amplified using kit-provided Pre-amplifier and Amplifier conjugated to alkaline phosphatase and incubated with a Fast Red substrate solution for 10 min at room temperature. Sections were then stained with hematoxylin, air-dried, and cover slipped.

### Data analysis

Telemetry data in the NSS files were extracted and further reduced into a validated MS Excel workbook for each NHP using Notocord-derived formula add-ins. Data reduction was done in 30-min intervals for temperature, 12-hr intervals for activity. Data collected for 4 days prior to virus exposure were used to generate a baseline dataset used for comparisons post challenge. 30-min baseline data points were calculated by averaging the time-matched values from each baseline day. In addition to the Notocord add-in functions, the MS Excel workbook contained Excel links and formulas, such as average and standard deviation (SD), used to further process the Notocord-derived data and contained logical functions that highlighted study values that were 3 SD above or below their concomitant baseline values. Fever was defined as body temperature >1.5°C above time-matched baseline for longer than 2 hours. Hyperpyrexia was defined as body temperature >3.0°C above time-matched baseline for longer than 2 hours. Severe hypothermia was defined as body temperature >2.0°C below time-matched baseline for 30 minutes. These highlighted study values were used to populate graphs with body temperature values that were significantly changed from baseline. Telemetry data collected from the challenge period (time period from the virus exposure to the endpoint when the animal is removed from cage for euthanasia) were used as challenge telemetry data.

### Statistics

All analyses were performed using GraphPad Prism 9. Data are presented as mean ± SEM. Statistical analysis of PRNT80 GMTs was performed using unpaired 2-tailed Student’s *t* test, with a *P* value of less than 0.05 considered significant. Statistical analysis of radiographic scores, fever-hr, maximum temperature change, and daily percent TEsig duration was performed using the Kruskal-Wallis test, with a *P* value of less than 0.05 considered significant.

### Study approval

Research was conducted under an IACUC-approved protocol in compliance with the Animal Welfare Act, PHS Policy, and other Federal statutes and regulations relating to animals and experiments involving animals.

The animal study was performed in ABSL3 containment facilities at USAMRIID. The facility where this research was conducted is accredited by the Association for Assessment and Accreditation of Laboratory Animal Care, International and adheres to principles stated in the Guide for the Care and Use of Laboratory Animals, National Research Council, 2011.

## Author Contributions

SLB, AJ, FR, JWH, KMG, TDM, AN, and MLMP conceptualized the project. SLB, AJ, FR, KMR, AMM, XZ, JWH, JWK, BK, KG, TDM, AN, and MLMP designed the study and developed the assays. SLB, CPS, AJ, FR, KMR, CJS, AMM, XZ, JWH, DD, OF, BK, JL, ST, TLC, JMS, JAJ, KB, HE, KLD, SRC, HB, PK, and KA performed the research. SLB, CPS, AJ, FR, KMR, CJS, AMM, XZ, JWH, DD, BK, ND, JL, ST, JAJ, KB, and KA analyzed the data. SLB, AJ, FR, AMM, XZ, and MLMP wrote the initial draft of the manuscript. SLB, CPS, KMR, CJS, AMM, XZ, and JWH designed the figures. SLB, CPS, AJ, FR, KMR, CJS, AMM, XZ, JWH, JWK, BK, ND, KB, KG, TDM, AN, and MLMP reviewed the manuscript and provided feedback and edits.

## Acknowledgments

The authors acknowledge the members of the Veterinary Medicine Division for assistance with all animal-related activities and study planning and execution, along with Aerobiology, Animal Clinical Pathology, and Telemetry for technical assistance and study execution. The authors also thank Kristan O’Brien for program and timeline management, assistance with study execution, and administrative support, Dr. Sara Johnston for insightful discussions, and Sarah Norris for statistical consultations. We also acknowledge Denise Danner and James Barth of Core Laboratory Services for their technical assistance with production of the virus stock and plaque assays. Funding for this effort was provided by the Military Infectious Diseases Research Program under project number 150155790.

## References

1. Hewitt JA, et al. ACTIVating Resources for the COVID-19 Pandemic: In Vivo Models for Vaccines and Therapeutics. Cell Host Microbe. 2020;28(5):646–659.

2. Krammer F. Correlates of protection from SARS-CoV-2 infection. Lancet. 2021.

3. Chen N, et al. Epidemiological and clinical characteristics of 99 cases of 2019 novel coronavirus pneumonia in Wuhan, China: a descriptive study. Lancet. 2020;395(10223):507–513.

4. Huang C, et al. Clinical features of patients infected with 2019 novel coronavirus in Wuhan, China. Lancet. 2020;395(10223):497–506.

5. Wang D, et al. Clinical Characteristics of 138 Hospitalized Patients With 2019 Novel Coronavirus-Infected Pneumonia in Wuhan, China. JAMA. 2020;323(11):1061–1069.

6. Zhou F, et al. Clinical course and risk factors for mortality of adult inpatients with COVID-19 in Wuhan, China: a retrospective cohort study. Lancet. 2020;395(10229):1054–1062.

7. Polak SB, et al. A systematic review of pathological findings in COVID-19: a pathophysiological timeline and possible mechanisms of disease progression. Mod Pathol. 2020;33(11):2128–2138.

8. Munster VJ, et al. Respiratory disease in rhesus macaques inoculated with SARS-CoV-2. Nature. 2020;585(7824):268–272.

9. Deng W, et al. Ocular conjunctival inoculation of SARS-CoV-2 can cause mild COVID-19 in rhesus macaques. Nat Commun. 2020;11(1):4400.

10. Salguero FJ, et al. Comparison of rhesus and cynomolgus macaques as an infection model for COVID-19. Nat Commun. 2021;12(1):1260.

11. Hartman AL, et al. SARS-CoV-2 infection of African green monkeys results in mild respiratory disease discernible by PET/CT imaging and shedding of infectious virus from both respiratory and gastrointestinal tracts. PLoS Pathog. 2020;16(9):e1008903.

12. Blair RV, et al. Acute Respiratory Distress in Aged, SARS-CoV-2-Infected African Green Monkeys but Not Rhesus Macaques. Am J Pathol. 2021;191(2):274–282.

13. Johnston SC, et al. Development of a coronavirus disease 2019 nonhuman primate model using airborne exposure. PLoS One. 2021;16(2):e0246366.

14. Centers for Disease Control and Prevention. Science Brief: SARS-CoV-2 and Potential Airborne Transmission. https://www.cdc.gov/coronavirus/2019-ncov/science/science-briefs/scientific-brief-sars-cov-2.html. Updated October 5, 2020. Accessed March 30, 2021.

15. The Lancet Respiratory Medicine. COVID-19 transmission-up in the air. Lancet Respir Med. 2020;8(12):1159.

16. Klompas M, et al. Airborne Transmission of SARS-CoV-2: Theoretical Considerations and Available Evidence. JAMA. 2020;324(5):441–442.

17. Fennelly KP. Particle sizes of infectious aerosols: implications for infection control. Lancet Respir Med. 2020;8(9):914–924.

18. Siegel JD, et al. 2007 Guideline for Isolation Precautions: Preventing Transmission of Infectious Agents in Health Care Settings. Am J Infect Control. 2007;35(10 Suppl 2):S65–164.

19. Pourbagheri-Sigaroodi A, et al. Laboratory findings in COVID-19 diagnosis and prognosis. Clin Chim Acta. 2020;510:475–482.

20. Salinas M, et al. Laboratory parameters in patients with COVID-19 on first emergency admission is different in non-survivors: albumin and lactate dehydrogenase as risk factors. J Clin Pathol. 2020.

21. Driscoll KE, et al. Intratracheal instillation as an exposure technique for the evaluation of respiratory tract toxicity: uses and limitations. Toxicol Sci. 2000;55(1):24–35.

22. Brown JH, et al. Influence of Particle Size upon the Retention of Particulate Matter in the Human Lung. Am J Public Health Nations Health. 1950;40(4):450–480.

23. Harcourt J, et al. Severe Acute Respiratory Syndrome Coronavirus 2 from Patient with Coronavirus Disease, United States. Emerg Infect Dis. 2020;26(6):1266–1273.

24. Wolfel R, et al. Virological assessment of hospitalized patients with COVID-2019. Nature. 2020;581(7809):465–469.

25. Corman VM, et al. Detection of 2019 novel coronavirus (2019-nCoV) by real-time RT-PCR. Euro Surveill. 2020;25(3).

26. Wrapp D, et al. Cryo-EM structure of the 2019-nCoV spike in the prefusion conformation. Science. 2020;367(6483):1260–1263.

27. Liu J, et al. Molecular detection of SARS-CoV-2 in formalin-fixed, paraffin-embedded specimens. JCI Insight. 2020;5(12).

