## Supplementary Material for "Aerosol Exposure of Cynomolgus Macaques to SARS-CoV-2 Results in More Severe Pathology than Existing Models"

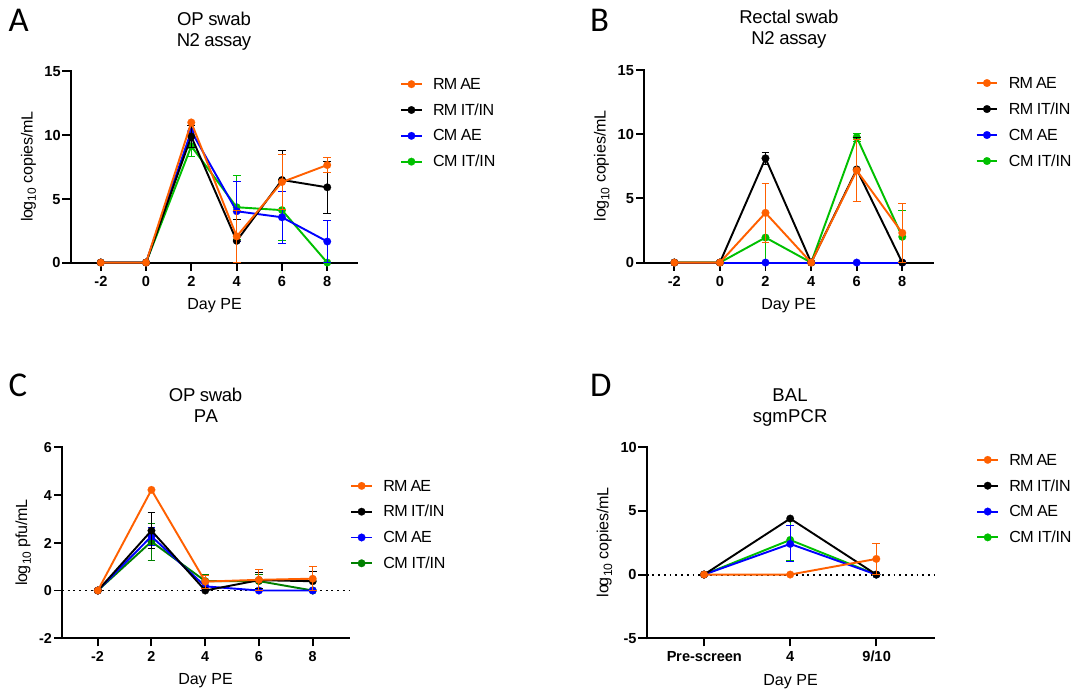

**Supplementary Figure 1**. Viremia in biosamples following infection with SARS-CoV-2. A-B. Viral RNA in OP (A) and rectal (B) swabs as detected by RT-PCR. C) Infectious virus in OP swabs as detected by plaque assay. D) Detection of subgenomic RNA in BAL samples using real-time RT-PCR. All data are shown as mean ± SEM.

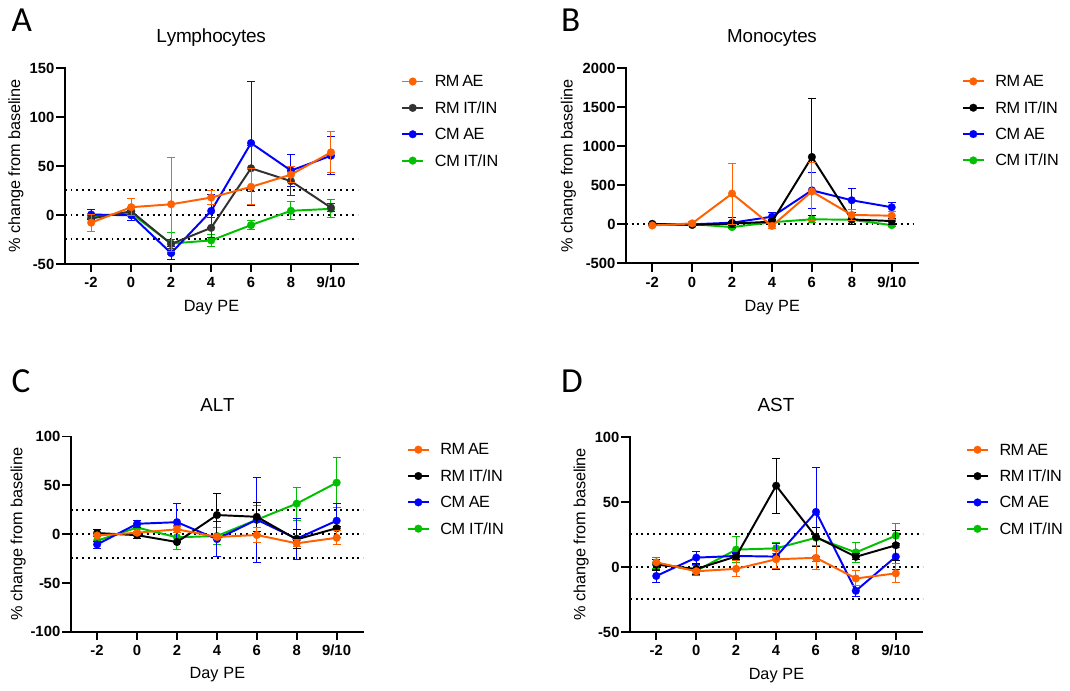

**Supplementary Figure 2**. Clinical pathology alterations following SARS-CoV-2 exposure. Percent change from baseline for A) lymphocytes, B) monocytes, C) ALT, and D) AST over the course of the study. The dashed line at y=0 represents the baseline, which is defined as the average of the results from Days -2 and 0. Values greater than or equal to a 25% change from baseline (dashed lines in A, C, and D) were considered noteworthy. All data are shown as mean ± SEM.

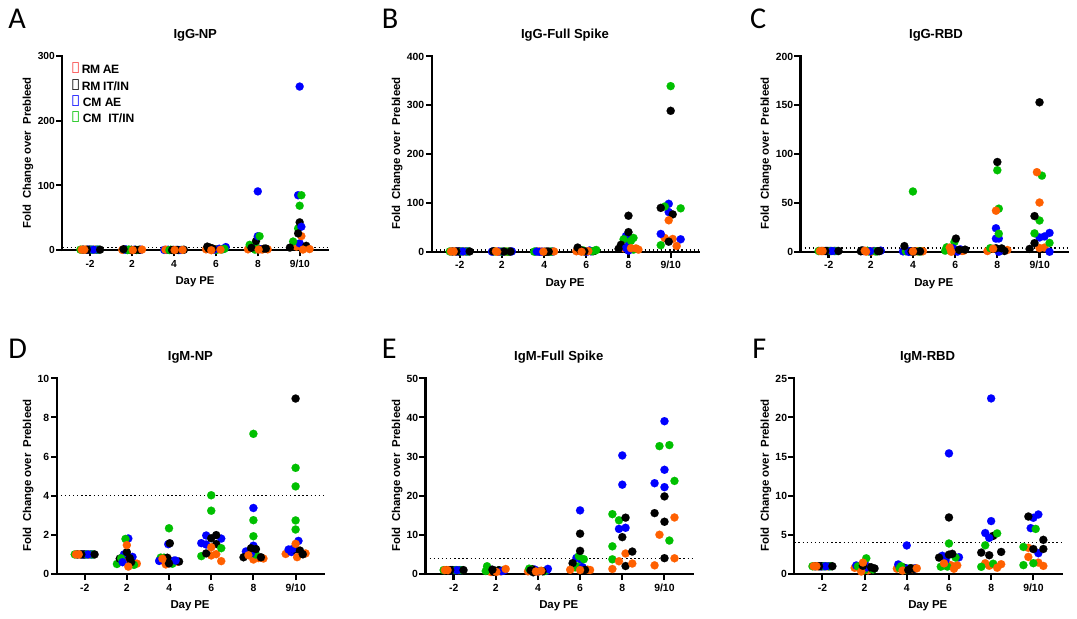

**Supplementary Figure 3.** Characterization of the IgG and IgM responses to SARS-CoV-2. A Magpix multiplex immunoassay was used to measure the IgG and IgM response to NP, full spike protein, and RBD of SARS-CoV-2 in serum samples. Results are shown as fold change over pre-bleed (Day -2). The dashed line represents the limit of detection of the assay.

**Supplementary Table 1**. Summary of histopathology findings in SARS-CoV-2 infected NHPs

| **Group** | **Animal** | **Pulmonary Lesions** | | | | | **ISH+** | **MGNCs^A^ present** |  | | | |
| --- | --- | --- | --- | --- | --- | --- | --- | --- | --- | --- | --- | --- |
|  |  | **Inflammation** | **Type II pneumocyte hyperplasia** | **Fibrin deposition alveolar lumen (strands)/ septa** | **Intra-alveolar fibrin/ fibrous aggregated deposition** | **Septal fibrous change** |  |  |  |  |  |  |
| **RM AE** | **RM AE 1** | **1** | **2** | **0** | **0** | **0** | **No** | **Yes** |  |  |  |  |
|  | **RM AE 2** | **1** | **1** | **0** | **0** | **0** | **No** | **No** |  | **Scoring Key** | | |
|  | **RM AE 3** | **1** | **1** | **0** | **0** | **0** | **Yes** | **Rare** |  | **0** |  | **None** |
|  | **RM AE 4** | **2** | **2** | **0** | **0** | **1** | **Yes** | **Rare** |  | **1** |  | **Minimal** |
| **RM IT/IN** | **RM IT/IN 1** | **1** | **1** | **0** | **0** | **0** | **Yes** | **No** |  | **2** |  | **Mild** |
|  | **RM IT/IN 2** | **1** | **1** | **0** | **0** | **1** | **No** | **Rare** |  | **3** |  | **Moderate** |
|  | **RM IT/IN 3** | **1** | **0** | **0** | **0** | **1** | **No** | **No** |  | **4** |  | **Marked** |
|  | **RM IT/IN 4** | **1** | **0** | **0** | **0** | **0** | **No** | **No** |  | **5** |  | **Severe** |
| **CM AE** | **CM AE 1** | **3** | **3** | **3** | **3** | **2** | **Yes** | **Yes** |  | | | |
|  | **CM AE 2** | **3** | **3** | **2** | **0** | **2** | **Yes** | **Yes** |  |  |  |  |
|  | **CM AE 3** | **3** | **3** | **2** | **0** | **2** | **Yes** | **Yes** |  |  |  |  |
|  | **CM AE 4** | **3** | **3** | **3** | **3** | **3** | **Yes** | **Yes** |  |  |  |  |
| **CM IT/IN** | **CM IT/IN 1** | **2** | **2** | **1** | **0** | **2** | **Yes** | **Yes** |  |  |  |  |
|  | **CM IT/IN 2** | **2** | **2** | **2** | **0** | **2** | **Yes** | **Yes** |  |  |  |  |
|  | **CM IT/IN 3** | **3** | **3** | **2** | **1** | **2** | **Yes** | **Yes** |  |  |  |  |
|  | **CM IT/IN 4** | **2** | **2** | **1** | **1** | **1** | **No** | **Yes** |  |  |  |  |

^A^ MGNCs = multinucleated giant cells

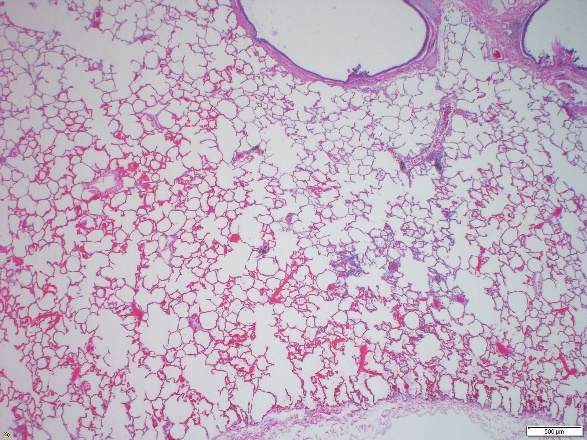

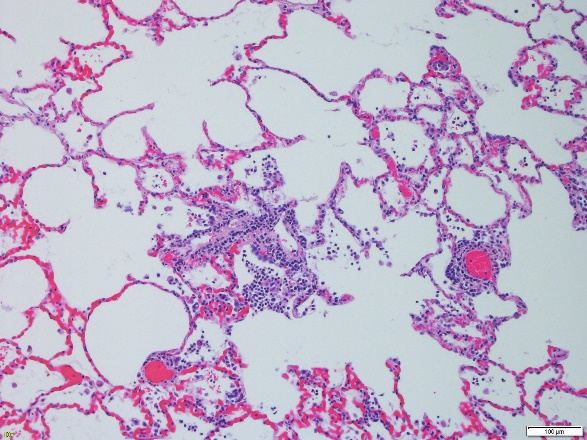

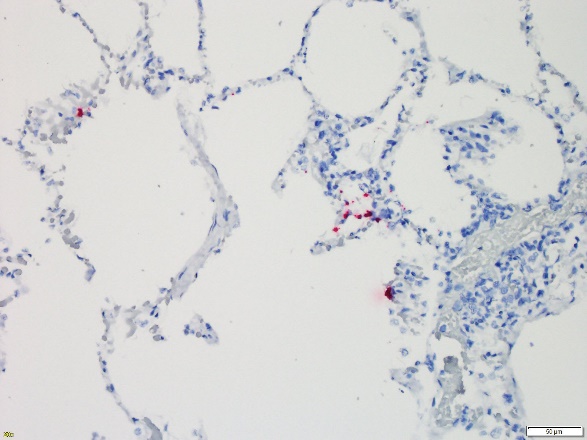

**Supplementary Figure 4**. Representative pathology in the RM AE group. Images are shown from RM AE 3. A. Lung, left cranial lobe, central: Multifocal minimal interstitial pneumonia, 2x, H&E. B. Lung, higher magnification of boxed area in A: Multifocal minimal inflammation surrounding vessels, expanding alveolar septa and extending into alveolar lumen with minimal type II pneumocyte hyperplasia, 10x, H&E. C. Lung: ISH positive in areas of inflammation, 20x, RNA probe for SARS-CoV-2.

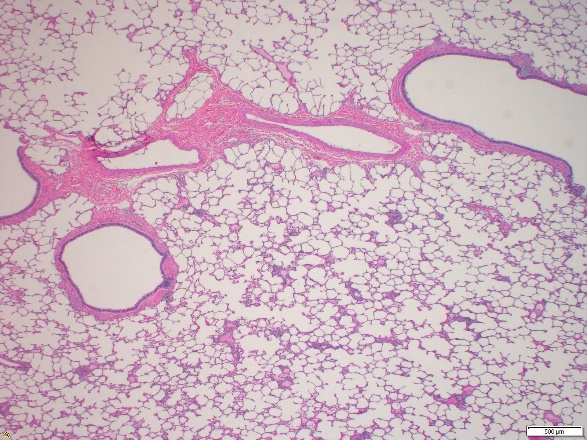

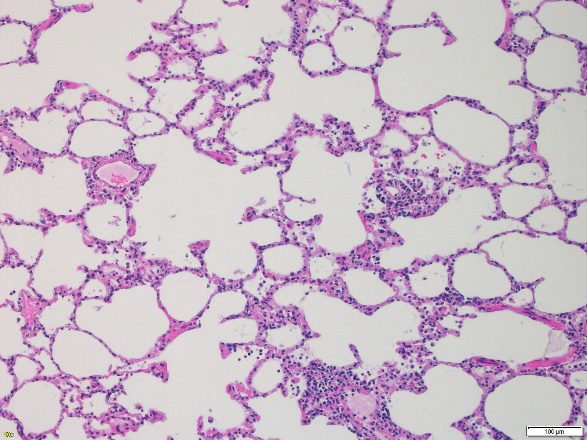

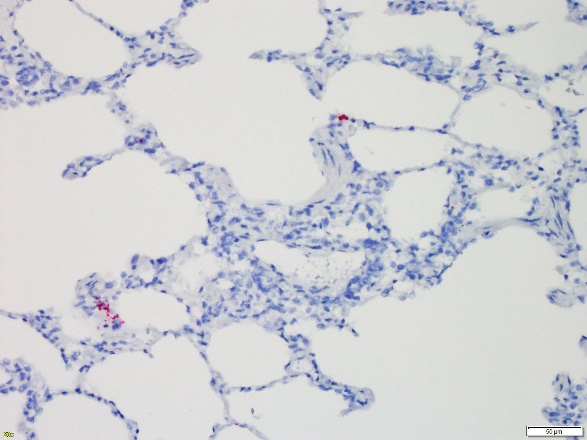

**Supplementary Figure 5**. Representative pathology in the RM IT/IN group. Images shown are from RM IT/IN 1. A. Lung, right caudal lobe, peripheral: Multifocal minimal interstitial pneumonia, 2x, H&E. B. Lung, higher magnification of boxed area in A: Multifocal minimal inflammation surrounding vessels, expanding alveolar septa and extending into alveolar lumen with minimal type II pneumocyte hyperplasia, 10x, H&E. C. Lung: ISH positive in areas of inflammation, 20x, RNA probe for SARS-CoV-2.

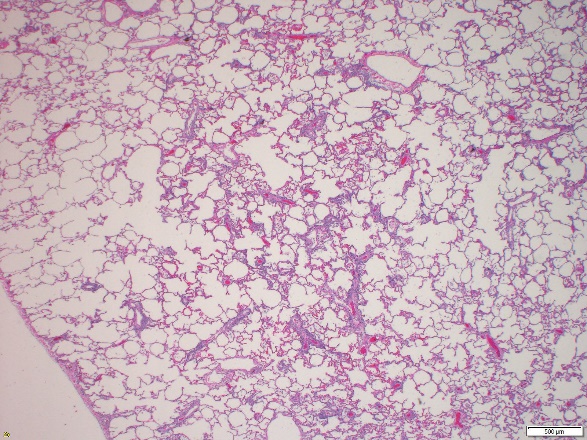

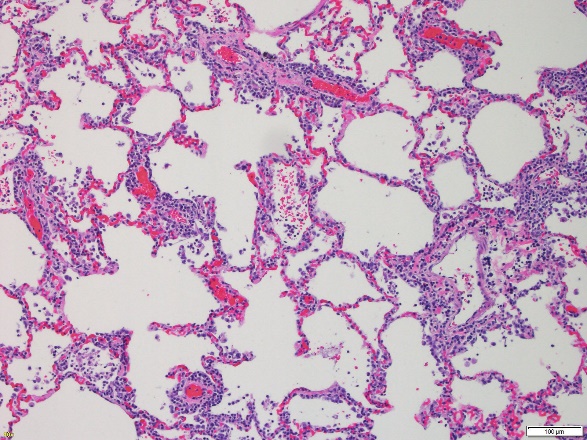

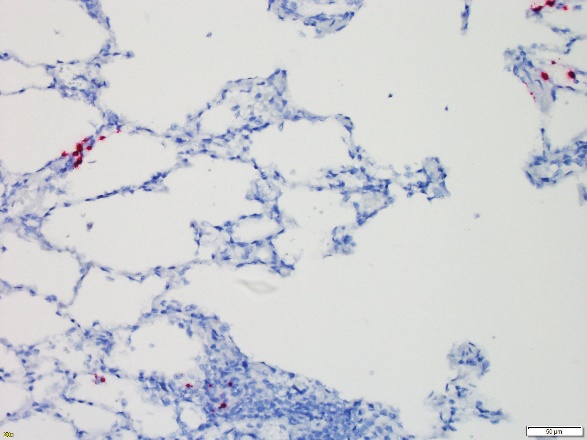

**Supplementary Figure 6**. Representative pathology in the CM IT/IN group. Images shown are from CM IT/IN 1. A. Lung, right caudal lobe, central: Multifocal mild interstitial pneumonia, 2x, H&E. B. Lung, higher magnification of boxed area in A: Multifocal mild inflammation surrounding vessels, expanding alveolar septa and extending into alveolar lumen with mild type II pneumocyte hyperplasia, 10x, H&E. C. Lung: ISH positive in areas of inflammation, 20x, RNA probe for SARS-CoV-2.
